# Actin bundle architecture and mechanics regulate myosin II force generation

**DOI:** 10.1101/2020.07.17.207415

**Authors:** Kimberly L. Weirich, Samantha Stam, Ed Munro, Margaret L. Gardel

## Abstract

The actin cytoskeleton is a soft, structural material that underlies biological processes such as cell division, motility, and cargo transport. The cross-linked actin filaments self-organize into a myriad of architectures, from disordered meshworks to ordered bundles, which are hypothesized to control the actomyosin force generation that regulates cell migration, shape, and adhesion. Here, we use fluorescence microscopy and simulations to investigate how actin bundle architectures with varying polarity, spacing, and rigidity impact myosin II dynamics and force generation. Microscopy reveals that mixed polarity bundles formed by rigid cross-linkers support slow, bidirectional myosin II filament motion, punctuated by periods of stalled motion. Simulations reveal that these locations of stalled myosin motion correspond to sustained, high forces in regions of balanced actin filament polarity. By contrast, mixed polarity bundles formed by compliant, large cross-linkers support fast, bidirectional motion with no traps. Simulations indicate that trap duration is directly related to force magnitude, and that the observed increased velocity corresponds to lower forces resulting from both the increased bundle compliance and filament spacing. Our results indicate that the properties of actin structures regulate the dynamics and magnitude of myosin II forces, highlighting the importance of architecture and mechanics in regulating forces in biological materials.

## Introduction

The actin cytoskeleton is an active biopolymer structure that underlies the mechanical behavior of cells, such as their ability to change shape and interact with their surroundings (1). This complex material is constructed from actin filaments (F-actin) that are intricately arranged into various architectures by proteins that cross-link F-actin and control F-actin nucleation and growth (2, 3). Cross-linkers bind to two F-actin, promoting the formation of networks and bundles with microstructures and mechanics that vary with cross-linker structure. For example, large, compliant cross-linkers, such as filamin, favor formation of networks with little angular constraint between filaments, that coexist primarily with fine cortical structures (4). By contrast, short, rigid cross-linkers such as α-actinin, fimbrin, and fascin form tightly packed, stiff bundles, capable of bearing loads and acting as cellular tracks, in the cell’s interior and protrusions (5, 6). Together, these and other proteins build actin architectures that control cell shape, mechanics, and intracellular transport (2, 3).

Embedded myosin motor proteins transform passively cross-linked actin architectures into active materials (7). Myosin motors undergo a mechanochemical cycle during which adenosine triphosphate (ATP) hydrolysis results in the motor binding to F-actin, exerting a force on the F-actin, and detaching from F-actin to restart the cycle. Myosin force generation was originally investigated in striated muscle, where actomyosin assemblies have well-defined sarcomeric organization (8). In sarcomeres, collections of approximately 500 myosin II motor heads polymerize into a bipolar filament, where the heads each exert step-like forces in a directed fashion towards the barbed end of F-actin. The microstructure of a sarcomere unit, where anchored F-actin is arranged with opposing polarity, is key to the collective myosin II forces resulting in a net contractile force (9). Myosin II force generation is also critical in nonmuscle cells, where F-actin assemblies lack well defined sarcomeric order, but instead have disordered polarity, polydisperse filament length, as well as various angular orientations and mechanical anchorings. The complex microstructures in non-muscle cells are likely to spatially regulate the activity of myosins and other proteins. For example, F-actin spacing in bundles influences protein binding and localization, while F-actin polarity and angle can direct transport (10-13). Unlike in sarcomeres where the F-actin arranged with opposing polarity supports contractile force generation, in filopodia F-actin are arranged with the same polarity, which may facilitate transport into cellular protrusions (2, 14). However, despite evidence that cytoskeletal architecture can impact protein localization and cargo motility generated by transport motor proteins (10, 13, 15-17), and network architecture is critical to myosin-driven contractility (18-20)the role of F-actin architecture in myosin II force generation remains an open question.

Here, we investigate the impact of F-actin architecture on myosin II-based forces in model actomyosin experiments reconstituted from purified proteins and through complementary agent-based simulations. Through fluorescence microscopy, we find that isolated filaments of skeletal muscle myosin II (myosin) move throughout space on cross-linked F-actin networks, but have bidirectional motion confined to micron-sized regions on F-actin bundles. To understand the origins of the confinement, we construct actin bundle architectures *in vitro* and *in silico* with different filament polarity and spacing. On mixed polarity bundles with small interfilament spacing, we find that myosin moves bidirectionally, at speeds nearly an order of magnitude below the gliding velocity, punctuated by periods in which motors are nearly stalled. By contrast, on mixed polarity bundles formed by cross-linkers which support large interfilament spacing and angular flexibility, myosin moves bidirectionally without stalling. Intriguingly, increasing the amount of cross-linking in these spaced, compliant bundles causes myosin motion to be similar to smaller-spaced, rigid bundles, with regions of confined, stalled motion. The simulations show that while the myosin is robustly confined to traps in balanced polarity regions with small interfilament spacing, the relationship between the force the myosin exerts on F-actin bundle and the filament microstructure is complex. Through simulations we can decouple the effects of cross-linker spacing and compliance. In order to capture the dynamics that we see in experimentally constructed bundles with large, compliant cross-linkers, it is necessary that cross-linkers impose a large spacing, while the low spring constants enhance the bidirectional motion. Our results show that myosin motion and forces are sensitive to bundle architecture, indicating that cytoskeletal microstructure and mechanics may be an important regulator of cellular force production.

## Methods

### Experimental assay

The sample chamber is a channel, formed by a thin (2 mm) Teflon gasket (McMaster Carr) pressed against a boroslicate coverslip (Fisherbrand). Glass and Teflon are rinsed with pure ethanol (200 proof; Decon Laboratories), then pure water (milliQ), and pure ethanol again, before drying with a stream of air. Glass is then exposed to UV/ozone (UVO cleaner; Jetlight) for at least 15 min, before immediately assembling the sample chamber and filling with vesicle buffer (140 mM NaCl, 8.5 mM Na_2_HPO_4_, 1.5 mM NaH_2_PO_4_, pH 7.5). The glass surface is passivated against protein adhesion by a supported lipid bilayer. After drying films of DOPC (1,2-dioleoyl-sn-glycero-3-phosphocholine, Avanti Polar Lipids) under filtered nitrogen gas, and resuspending in vesicle buffer, the suspension is extruded (200 nm and 50 nm pore membranes, Liposofast extruder, Avestin) to form unilamellar vesicles. Incubating the sample chamber with 1 mM vesicle suspension results in a complete supported lipid bilayer in <5 min. The sample solution is then exchanged with F-actin polymerization buffer (10 mM imidazole, 50 mM KCl, 0.2 mM EGTA pH 7.5, 300 uM ATP).

Adding monomeric actin (rabbit skeletal muscle purified from acetone powder (21), Pel-freeze Biologicals, Rogers, AR; 2.0 µM unlabeled and 0.64 µM labeled with the fluorophore tetramethylrhodamine-6-maleimide, Life Technologies) caused the polymerization of F-actin. A small amount of depletant (0.3 wt%, 15 centipoise methylcellulose, Sigma) crowds the F-actin to a thin layer near the surface. After 30 min of polymerization, protein cross-linker is added to bundle the F-actin. Fascin (human, purified from bacteria (22), courtesy of D. Kovar lab), Fimbrin (pombe (23), courtesy of D. Kovar lab), a-Actinin (human, purified from insect cells (24), courtesy of D. Kovar lab), or filamin (purified from chicken gizzard using protocol modified from ref. (25)) added between 0.1 and 10 mol% to cross-link the F-actin. The cross-linked bundles coarsen for the next ∼30 min. After the network reaches a steady state, where coarsening is no longer visible, prepolymerized filaments of skeletal muscle myosin II and ATP is gently mixed in the chamber, such that the final concentrations were 3.8 pM myosin and 2.3 mM ATP. Myosin II filaments are prepolymerized by adding monomeric myosin II (20 nM, chicken skeletal muscle, fluorescently labeled with Alexa 647, Life Technologies) to an actin free sample solution and allowing to incubate for 10 min.

Oxygen scavenging system (50 µM glucose, 0.5 vol% β-mercaptoethanol, glucose oxidase, and catalase) minimizes photobleaching. The samples are imaged with an inverted microscope (Nikon Eclipse Ti-PFS) equipped with a spinning disk confocal head (CSUX, Yokogawa), 561 nm and 647 nm laser lines, 60×/1.49 NA oil immersion objective (Zeiss), and a CCD camera (Coolsnap HQ2, Photometrics). Images, collected at 1.5 s intervals, began ∼10 min after myosin is added to sample.

### Image analysis

The frame to frame velocity of myosin II puncta is obtained through using FSM center to locate and track myosin. Thresholded images of networks are obtained via imageJ. Using the bundle threshold as a mask via Matlab, velocities are selected that correspond to motors localized to bundles.

Maximal intensity projections are obtained through the built-in Max Projection function of ImageJ, collapsing 300 s of data at 1.5 s intervals onto a single plane. Regions of myosin localization along bundles appear as bright linear regions in the maximal intensity projection. A line is drawn along the bundle that extends through these bright regions identified in the maximal intensity projection. Then, the intensity along the line is plotted vertically through each timepoint in the image sequence using the built-in ImageJ function, Reslice.

### Simulation Methods

Agent-based simulations were conducted using software we have previously described (26) that has been benchmarked to reproduce the gliding speed and force-velocity curves of myosin II using relevant experimentally determined single-molecule parameters. F-Actin is represented as a rigid rod, actin cross-linkers are represented as flexible springs, and myosin filaments are represented as rigid rods with extruding elastic elements capped by F-actin binding sites that represent the motor domains. Each of these elastic elements have a spring constant describing resistance to motion parallel to a F-actin, *K*_x-bridge, par_, as well as a weaker spring constant for perpendicular motion, *K*_x-bridge, perp_ that prevents bound F-actin from drifting. Motor domains are initially unbound at the beginning of a simulation and then stochastically bind and unbind via the Gillespie algorithm with rates *k*_on_ and *k*_off_ (*F*) respectively. The unbinding rate *k*_off_ (*F*) follows the experimentally determined form (27):

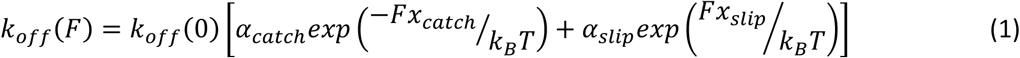

where the force *F* is positive for a load resisting the myosin step, *k*_B_ is Boltzmann’s constant, *T* is temperature, *x*_catch_ and *x*_slip_ are characteristic bond lengths, and α_catch_ and α_slip_ are constant prefactors. As a result of this form, the myosin bonds are catch bonds in which opposing forces slow the unbinding rate and assisting forces increase the unbinding rate provided that the force is below a threshold magnitude. Above this magnitude, the bond is a conventional slip bond where opposing forces increase unbinding. F-actin cross-linkers are bound at the beginning of the simulations and remain bound throughout the simulation. Binding sites for myosin or cross-linkers along the F-actin are spaced at a distance of *s*_binding site_ = 2.7 nm and myosin motors are spaced along their respective F-actin at a distance of *s*_motor_ = 5 nm. The values of all parameters are given in Table S1.

The myosin step is a result of the elastic motor domains binding actin in a prestrained state where the elastic element parallel to the actin filament is stretched by a distance *d*_step_. Subsequent relaxation of the elastic element causes the relative motion between the F-actin and myosin filaments. Motion of each filaments proceed by numerically solving its equation of motion:

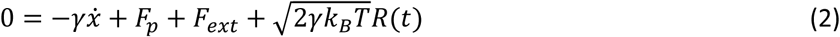

where γ is the drag coefficient in a medium with a viscosity of 0.1 N·s/m^2^, *F*_p_ is the force exerted by bound proteins, *F*_ext_ is an external load that is zero for myosin filaments and is imposed by anchoring springs for F-actin, *x* is the spatial coordinate, and *R*(t) is a random number drawn from a Gaussian distribution to yield the thermal force in final term of equation (2).

| Name | Description | Value | Reference |
| --- | --- | --- | --- |
| $N_{\text{heads}}$ | number of heads on one side of myosin filament | variable; 2-300 | - |
| $k_{\text{on}}$ | binding rate | $10 \text{ s}^{-1}$ | (28) |
| $k_{\text{off}}(\mathbf{0})$ | unloaded unbinding rate | $500 \text{ s}^{-1}$ | (29) |
| $\alpha_{\text{catch}}$ | See Eqn (1) | 0.92 | (27) |
| $\alpha_{\text{slip}}$ | See Eqn (1) | 0.08 | (27) |
| $x_{\text{catch}}$ | See Eqn (1) | 2.5 nm | (27) |
| $x_{\text{slip}}$ | See Eqn (1) | 0.4 nm | (27) |
| $k_{\text{off}}(\mathbf{F})$ | force dependent unbinding rate | See Eqn (1) | (27) |
| $d_{\text{step}}$ | step size | 5.5 nm | (30) |
| $K_{\text{x-bridge, par}}$ | myosin cross-bridge stiffness parallel to F-actin | 0.7 pN/nm | (30) |
| $K_{\text{xbridge, perp}}$ | myosin cross-bridge stiffness perpendicular to F-actin | 0.2 pN/nm | - |
| $k$ | stiffness of F-actin cross-linkers and springs that anchor F-actin | variable; 0.001-10 pN/nm | (31, 32) |
| $s_{\text{binding site}}$ | spacing of binding sites for motors or cross-linkers along F-actin | 2.7 nm | - |
| $s_{\text{motor}}$ | spacing between myosin motors | 5 nm | - |
| $s$ | spacing between F-actin | variable; 10-200 nm | - |

## Results

To investigate the impact of actin architecture on myosin II dynamics and force generation, we use confocal microscopy to image single skeletal muscle myosin II filaments on F-actin assemblies reconstituted *in vitro*. We polymerize F-actin (2.6 μM monomeric actin), crowd it into a thin layer near the surface of a passivated coverslip with a depletion agent and cross-link the F-actin with the physiological cross-linker, α-actinin. At low concentration (0.1 mol%) α-actinin cross-links the F-actin into networks; as the α-actinin concentration increases to 10 mol%, F-actin form bundles within the network (Fig. 1A & B). Once the network forms, we add pre-formed myosin II filaments which appear as isolated puncta (Fig. 1B, white puncta). The number density of myosin puncta on the network is sufficiently low (∼5×10^−3^ myosin puncta/µm^2^) that they do not generate enough force to deform the network (33, 34).

**Figure 1:**
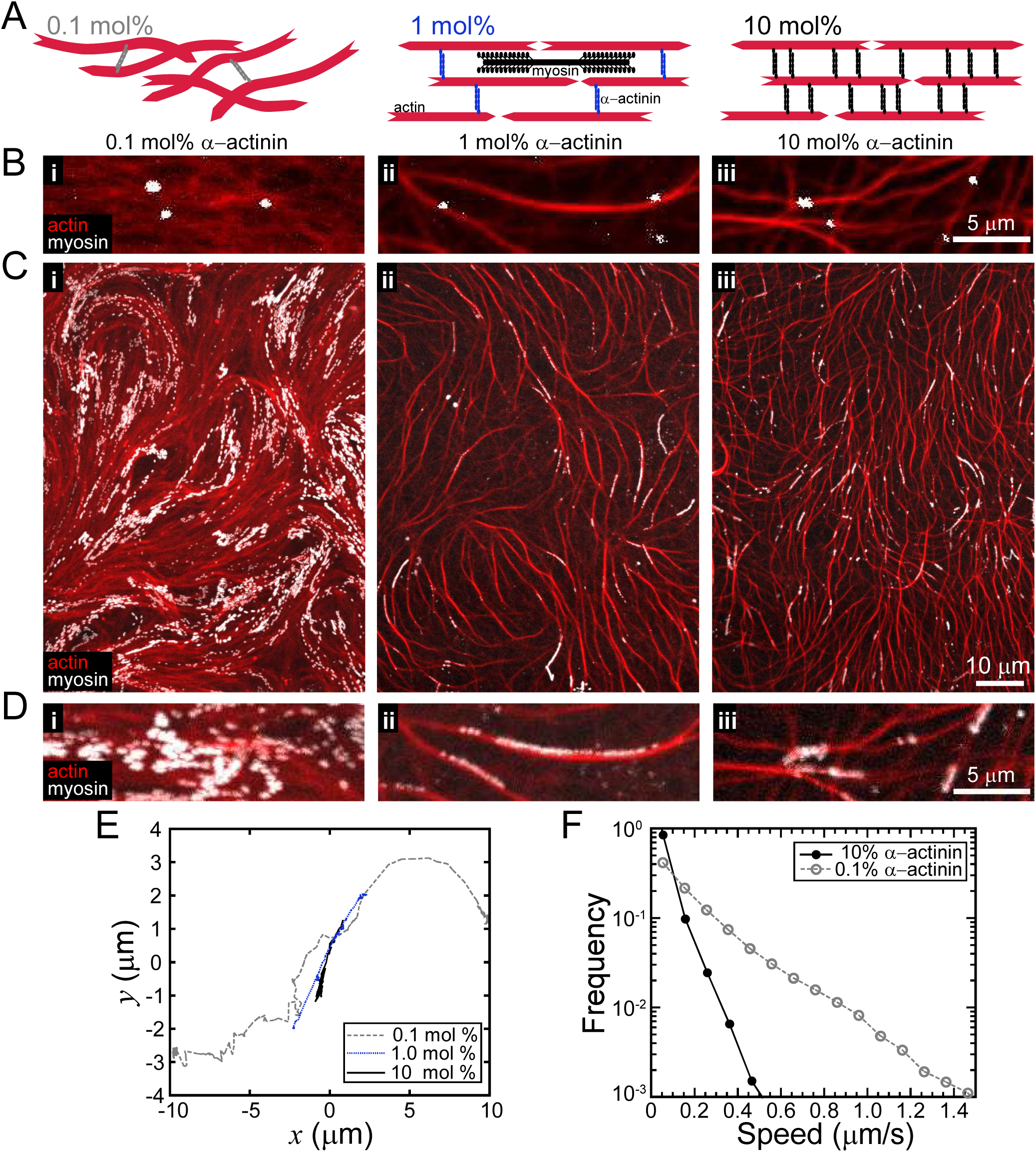
F-actin network architecture regulates myosin II motility. (A) Cross-linker constructs F-actin into architectures such as networks and bundles. (B) Fluorescence microscope images of F-actin (red) cross-linked by α-actinin with sparse myosin II filaments (white puncta). Network architecture is modulated by α-actinin concentration: networks of F-actin (left), networks with slight bundling (middle), and networks of bundles (right) form by cross-linking F-actin with 0.1 mol%, 1 mol%, and 10 mol% α-actinin, respectively. Isolated myosin II filaments (white puncta) localize to the networks. (C) Maximal intensity projection over 300 s of myosin II puncta dynamics on networks. (D) Zoomed in region of maximal intensity projection of myosin II on F-actin architectures in part B. (E) Representative trajectories of myosin II on networks with increasing amounts of α-actinin, transitioning between networks (0.1 mol%) and networks of bundles (1 mol%, 10 mol%). Motors on sparsely cross-linked networks explore more space, while motors on higher cross-linked bundled networks are restricted to linear movement along a particular bundle. (F) Distribution of myosin II instantaneous velocities, indicating that motor speed is reduced on higher cross-linked networks.

On sparsely cross-linked (0.1 mol%) F-actin, myosin II puncta are highly motile. Summing the myosin intensity over a period of 5 minutes and projecting on a single image reveals regions of extended tracks sparsely distributed over a large area (Fig. 1Ci & 1Di). By contrast, in the corresponding projections on F-actin bundles cross-linked by 1-10 mol% α-actinin, myosin puncta localize to dense, micron-sized linear regions, along individual F-actin bundles (Fig. 1Cii, 1Dii, 1 mol%; Fig. 1Ciii Fig. 1Diii, 10 mol%). To quantify differences in myosin speed, we tracked single myosin puncta (Fig. 1E). The probability distribution reveals that that on α-actinin cross-linked bundles, myosin have instantaneous speeds of <50 nm/s ∼80% of the time (Fig. 1F, black closed circles). By contrast, on the cross-linked actin networks myosin have speeds >50 nm/s and60% of the time (Fig. 1F, grey open circles), indicating that the amount of α-actinin-mediated bundling influences myosin puncta speed.

### F-actin polarity regulates myosin II filament velocity on rigid bundles

To elucidate the marked influence F-actin bundling has on myosin dynamics, we construct different F-actin bundle architectures from physiological cross-linkers. The most basic F-actin bundle architecture is a unipolar bundle in which the F-actin are all oriented in the same direction (Fig. 2A). To construct bundles with this architecture, we cross-link F-actin with fascin, a globular protein that mediates unipolar F-actin bundle formation with interfilament spacing <10 nm (Fig. 2Bi) (6, 10). Intensity projections over 5 min, reveals myosin puncta at regular intervals along the fascin cross-linked bundle, indicating uniform myosin speed (Fig. 2Ci). From a kymograph obtained along a bundle, we find that myosin puncta move persistently in one direction with a constant velocity (Fig. 2Di, left, Movie S4). Over 104 tracks, we find that ∼90% of myosin move persistently and ∼100% unidirectionally (Fig. 2E & F, blue), consistent with previous reports of single motor dynamics on unipolar bundles (16, 35). Quantifying the frame-to frame instantaneous velocity, indicates myosin has an average velocity of ∼2.5 μm/s on fascin bundles (Fig. 2G, blue solid), consistent with the unloaded velocity of skeletal muscle myosin II reported from gliding filament assays of single F-actin (36, 37). This indicates that unipolar bundles do not impact the gliding of myosin II filaments, suggesting that myosin puncta exert low forces on the unipolar bundles.

**Figure 2:**
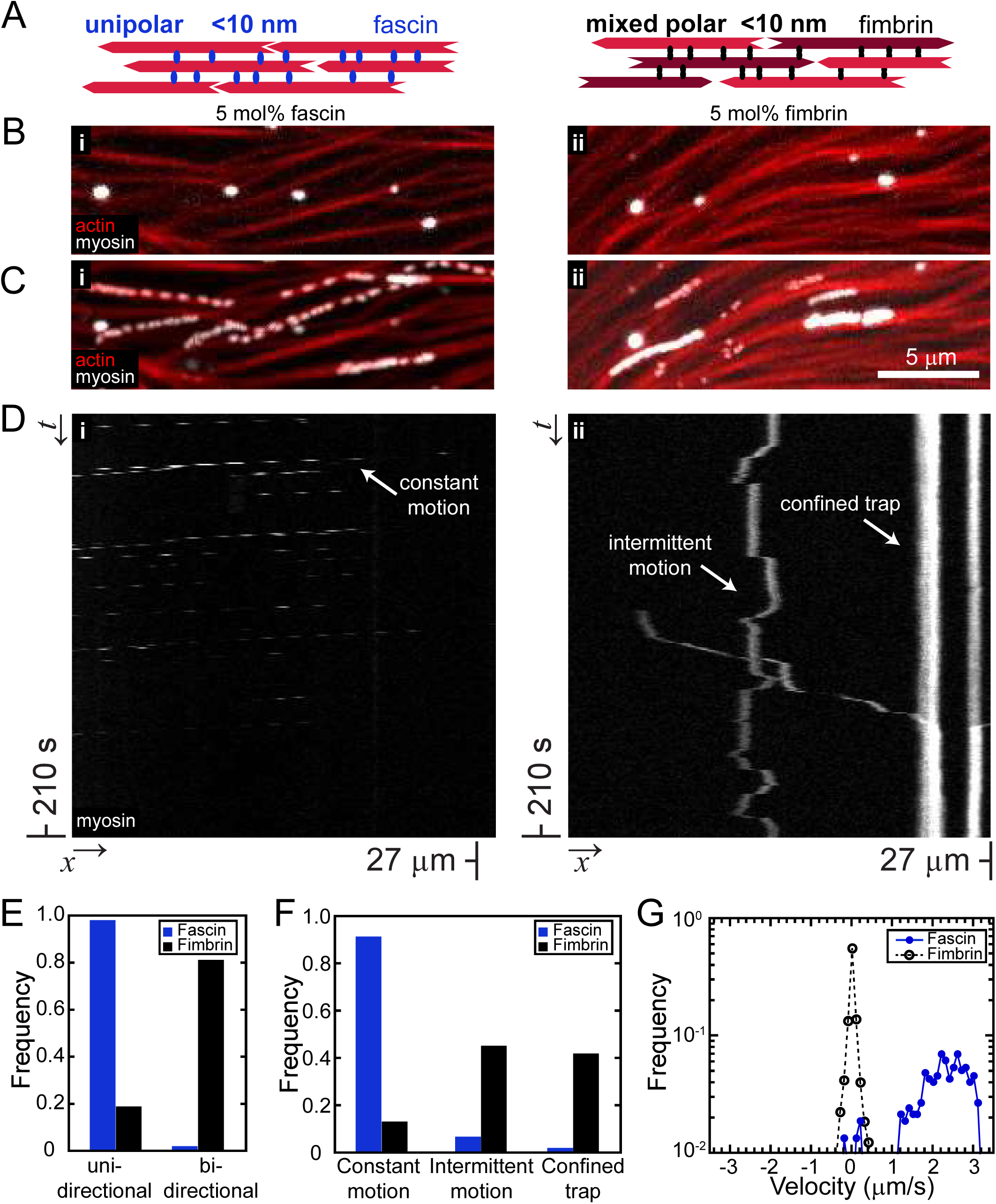
F-actin bundle polarity regulates myosin II dynamics. (A) Cartoon of tight unipolar bundles formed by fascin, where F-actin are oriented in the same direction (left) and mixed polarity bundles formed by fimbrin, where F-actin are oriented in both directions (right). (B) F-actin bundles (red) cross-linked by 5 mol% of the unipolar crosslinker, fascin (left), and the mixed polarity cross-linker, fimbrin (right), with isolated myosin II filaments (white puncta). (C) Maximal projection of myosin II (white) over 5 min on fascin-bundled F-actin (left, red) and fimbrin-bundled F-actin (right, red). On fascin-bundled F-actin, myosin appears as separated puncta distributed evenly along the bundles, whereas on fimbrin-bundled F-acitn, myosin localizes to dense regions of a few microns. (D) Kymograph of myosin II motor on fascin-bundled (left) and fimbrin-bundled F-actin (right). Myosin move along parallel diagonal lines, indicating myosin that bind to the bundle at different times and move with a similar constant velocity in one direction along the bundle. In contrast, on fimbrin bundles, myosin undergoes periods of motion, with both positive and negative slopes indicating bidirectional motion, interspersed with periods of traps (vertical lines). (E) Proportion of myosin moving along a bundle in one or two directions, in a single processive run. (F) Summary of the frequency of myosin dynamics—intermittent motion and pauses, constant motion, or sustained pauses—from kymographs represented in (D). Each myosin (5 mol% fascin: 104 and 5 mol% fimbrin: 153 myosin) was observed over 300 s periods along a bundle section. (G) Myosin II velocity distribution on fascin bundles (left), indicates unidirectional motion with magnitude similar to unloaded gliding speed. In contrast, myosin II velocity distribution on fimbrin bundles (right) is symmetric about zero, indicating bidirectional motion, with a peak at zero, indicating strong trapping.

We next constructed bundles from cross-linkers that do not constrain the F-actin polarity within bundles. To change polarity without influencing bundle mechanics, we used fimbrin (5 mol%, Fig. 2Aii, 2Bii, & Movie S5), a cross-linker that maintains F-actin spacing similar to fascin (10, 38). Projections of the myosin intensity over 5 min appear as dense linear tracks along the bundle with limited extent (Fig. 2Cii, white), indicating myosin II puncta have slow velocities and are confined to isolated regions on mixed polarity bundles. Kymographs of the myosin trajectory along the bundle reveal diagonal lines, indicating periods of directed motion, punctuated by vertical lines, indicating periods of immotility, or traps (Fig. 2Dii). From inspecting 153 tracks, we find that ∼40% of myosin are trapped, while ∼45% of myosin move intermittently, and only ∼15% move continuously (Fig. 2F, black). From the 85 independent tracks of motile myosin, we find both positively and negatively sloped lines in ∼80% of the tracks (Fig. 2E, black), indicating bidirectional motion along the bundle, consistent with reports of single motor dynamics on mixed polarity bundles (16). Additionally, the kymograph lines have larger slopes on mixed polarity bundles than on the unipolar bundles, indicating slower instantaneous velocities (Fig. 2D). Quantifying the instantaneous velocity reveals that motile fraction of myosin puncta have speeds ∼300 nm/s, nearly an order of magnitude slower than those observed on unipolar bundles (Fig. 2G, black open).

### Traps are regions of balanced polarity, where myosin has sustained, high forces

Decreased myosin puncta speed and trapped periods on mixed polarity bundles could result from multiple physical mechanisms, such as direction switching, tug-of-war, or kinetic trapping (16, 34). Using a computational model, we investigate the microscopic mechanisms by which F-actin bundle architecture impacts myosin filament dynamics *in silico* (26). In each agent-based simulation, we place a bipolar myosin filament in the center of a F-actin bundle (Fig. 3A). The bipolar myosin filament is composed of two parallel arrays, connected by a central region. The individual motors are represented by a simple form of the swinging cross-bridge model and benchmarked to skeletal muscle myosin (26). The number of myosin motor domains, *N*_heads_, is equal on each end of the myosin filament. We construct bundles by anchoring individual F-actin via springs of stiffness, *k*, attached at both ends. Cross-linkers are represented by springs, which also have stiffness, *k*, and rest length, *s*, that sets the F-actin spacing in a bundle. The polarity, *p*, of a bundle is defined as the proportion of the F-actin oriented with barbed ends in the positive direction, *N*_+_ /(*N*_+_ + *N*_-_). To vary *p*, we hold the number of F-actin with barbed ends in the positive direction, *N*_+_, constant, while varying the number of F-actin with barbed ends in the opposing direction, *N*_-_.

**Figure 3:**
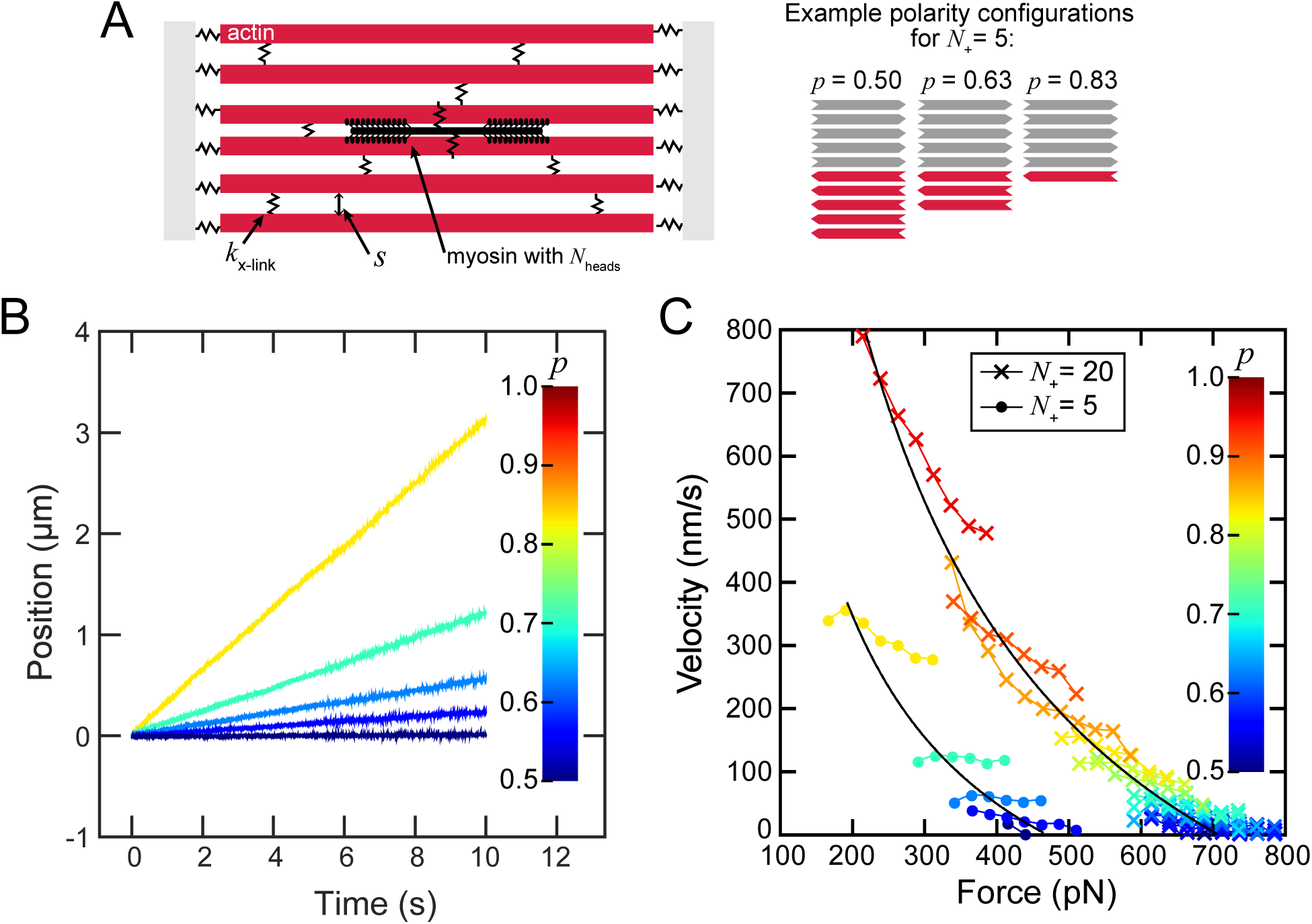
Regions of mixed polarity trap myosin filaments and maximize force output (A) Simulation schematic. A myosin filament is positioned at the center of an actin bundle, where actin filaments are oriented with barbed ends to the left or right. The highlighted myosin motor domains each are represented by springs that bind and unbind to F-actin with rates *k*_on_ and *k*_off_ respectively. The spring stretches upon binding, then relaxes, resulting in the myosin moving relative to the F-actin. The F-actin are attached to springs with stiffness, *k*, anchoring them in place and springs with stiffness, *k*, representing interfilament cross-links. (B) Trajectories of myosin filaments (*k* = 5 pN/nm) with varying actin polarity. The number of F-actin with barbed ends to the right, *N*_+_ is held constant at 5 while *N*_-_ is varied between 1 and 5. At larger values of *N*_-_ indicated by darker blue, myosin filaments are increasingly trapped in place and undergo small changes in direction. (C) Force-velocity curves in which the instantaneous velocity of the myosin filament is plotted against the instantaneous force stored in the myosin springs. Each reported point represents the average force, binned in increments of 25 pN. Blue curves include simulations where *N*_+_ is held constant and *N*_-_ is varied from 1 to 5 with darker shades indicating larger numbers. Red curves show equivalent data where *N*_+_ is equal to 20 while *N*_-_ is varied from 1 to 20. The value of *N*_heads_ is 250 for all curves. Each curve contains data from 20 independent 10 s simulations.

We systematically vary the polarity of a rigidly cross-linked (*k* = 5 pN/nm), tightly spaced (*s* = 0 nm), F-actin bundle. At this rigidity and spacing, the deformation of the F-actin springs and cross-linkers is negligible, resulting in an effectively rigid bundle. For *p* = 1, a completely unipolar bundle which corresponds to the experimental fascin-bundled F-actin, the myosin travels micrometers over a period of seconds in one direction, towards the F-actin barbed ends, consistent with experiments (Fig. 3B). As *p* decreases, the total myosin filament displacement also decreases, and the corresponding decreased velocity reflects an increasing resistance from motor binding to F-actin with opposing orientation. When *p* = 0.5, myosin filaments move only a few nanometers, similar to the confined motion we experimentally observe on mixed polarity bundles.

With this simulation, we can interrogate how the instantaneous force exerted by a myosin filament depends on the F-actin bundle polarity. We find the force generated on bundles with the most uniform polarity is lowest (Fig. 3C, red “x”), whereas the force exerted on bundles with mixed polarity is highest (Fig. 3C, blue “x”). Inspection of the forces exerted by the myosin filament on individual F-actin reveals that myosin binding to opposing F-actin in the bundle, *N-*, resist the myosin movement on F-actin oriented in the positive direction, resulting in the buildup of a force dipole. When the forces in the positive direction are not fully balanced by those in the opposing direction, processive motor motion in the positive direction occurs. This indicates that the forces a myosin filament exerts on the F-actin oriented in the positive direction, *N*_+_, are effectively determined by the motor interactions with F-actin of opposing polarity, *N-*. By plotting the average force and average myosin velocity measurements of bundles over a range of polarities, we see that these fall on a curve consistent with the known force-velocity relationship of myosin II which is encoded by the model (26) (Fig. 3C, black lines). The range of forces for each polarity reflect the stochasticity of the system and fluctuations in number of binding interactions. As the polarity becomes more mixed, the force approaches a maximum, determined by the stall force of each motor head and by the total number of motor interactions with F-actin. Consequently, reducing the number of F-actin in the bundle to 5 results in a similar force-velocity relationship, but with nearly half of the maximal force generated in a bundle with 20 F-actin (Fig. 3C, circles).

While experimental control of the myosin filament size (*N*_heads_) or the number of F-actin within a bundle is complex, we can vary the number of motor heads in simulations to gain further insight into the microscope mechanisms of force production. For mixed polarity bundles (*p* = 0.5, *N*_+_ = 3), myosin filaments with *N*_heads_ <40 have short runs with frequent direction switching (Fig. 4A, black). From the mean squared displacement as a function of delay time, we find a scaling exponent near one, indicating that the myosin filament motion is effectively diffusive, with an effective diffusion coefficient ∼0.1-0.2 μm^2^/s (Figure 4B & 4C). This is consistent with dynamics that arise from stochastic effects, with significant force imbalances leading to large displacements (26, 39). As *N*_heads_ increases > 40, the myosin movement is reduced (Fig. 4A, red). The motion remains diffusive, but the diffusion coefficient decreases to nearly zero (Fig. 4B & 4C). Thus, for *p*=0.5, the myosin filament motion is diffusive over all *N*_heads,_ but the diffusion constant decreases with increasing *N*_heads_ (Fig. 4C) (40). Above a critical value of *N*_heads_, the force—which is the product of *N*_heads_ and the duty ratio, or fraction of time that individual motors spend bound to F-actin (26)— increases proportionally with *N*_heads_ (Fig. 4D). In cells, different isoforms of myosin II forms filaments, composed of multiple motor heads, ranging from several to several hundred motor heads. This data suggests that as the number of heads increases, the probability of at least one bound motor engaging to F-actin of both polarity increases, a prerequisite to building force. As this probability increases, the overall diffusivity of the motor on the mixed polarity bundle decreases, resulting in minimal displacement and maximal force build up. Together, these data highlight the impact of the number of motor heads in potential of collections of motors to generate forces.

**Figure 4:**
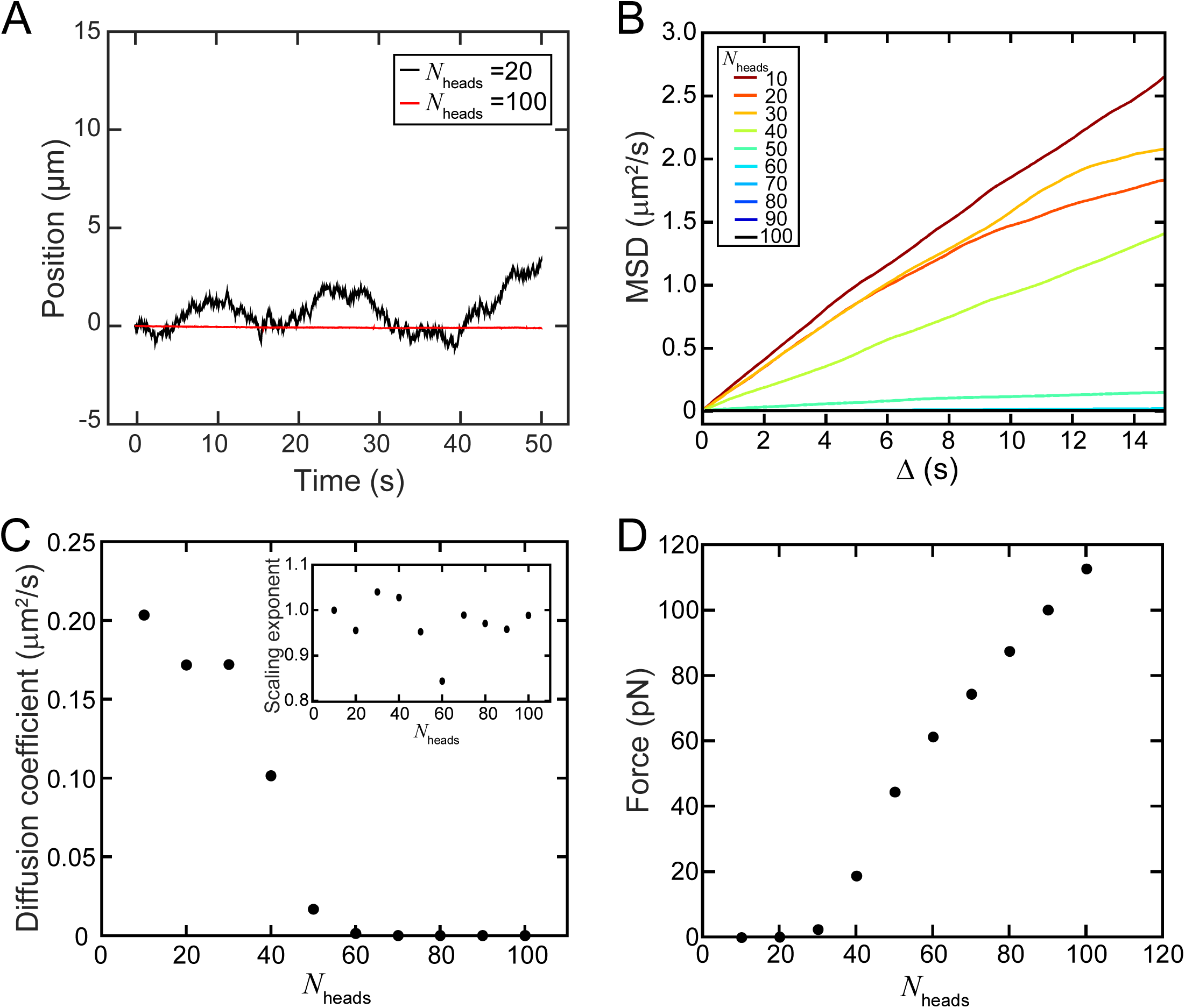
Number of myosin heads influences myosin dynamics and force generation. (A) Trajectories on *s*=10, *p*=0.5 bundles for *N*_heads_ = 20 and *N*_heads_ = 250. (B) Mean squared displacement as a function of delay time for bundles with *s*=10 nm and *k*=10 pN/nm. (C) Myosin filament diffusion coefficient, *D*, as a function of the number of heads, extracted from a fit to the data in (C), *MSD* = *De*^*β*^. Inset is the scaling exponent, β as a function of the *N*_heads_. (D) Force as a function of the number of myosin heads for myosin on a 10 nm spaced bundle, *N*_+_ = *N*___ = 3 with rigid cross-links.

### Myosin trapping is abrogated on filamin bundles and robust on a-actinin bundles

In addition to F-actin polarity, both the interfilament spacing and bundle compliance can be varied when constructing F-actin bundles with different cross-linkers. We hypothesize that changing the interfilament spacing potentially impacts the number of accessible F-actin binding sites for a motor complex bound to a bundle, while changes in the local bundle compliance could influence the force-dependent motor binding affinity (26, 40). Thus, we expect that the interfilament spacing and bundle compliance effect the number of motor heads bound, impacting the force generation potential of a given bundle architecture.

To investigate this, we experimentally constructed bundles formed with the cross-linkers a-actinin and filamin. These cross-linkers form mixed polarity bundles, similar to fimbrin, but form bundles with different interfilament spacing and compliance (4, 10, 41, 42). Cross-linking with α-actinin forms bundles with ∼35 nm interfilament spacing (Fig. 5A & 5Bi) (41). Projections of myosin intensity on a-actinin cross-linked bundles over a 5 min period appear as dense tracks along micron-sized regions of bundles (Fig. 5Ci). Despite different spacing, kymographs reveal regions of bidirectional motion and pauses, indicating myosin has similar motion as it does on fimbrin cross-linked bundles (Fig. 5Di, Movie S3). Over 217 individual myosin tracks, ∼60% of myosin move intermittently, similar to what we observe on fimbrin bundles (Fig. 5E, solid blue, closed circles). However, in contrast to fimbrin bundles, on α-actinin cross-linked bundles myosin are only continuously confined to a trap over a 5 min period in ∼8 % of the tracks, while they continuously move in ∼35 % of the cases (Fig. 5E, blue).

**Figure 5:**
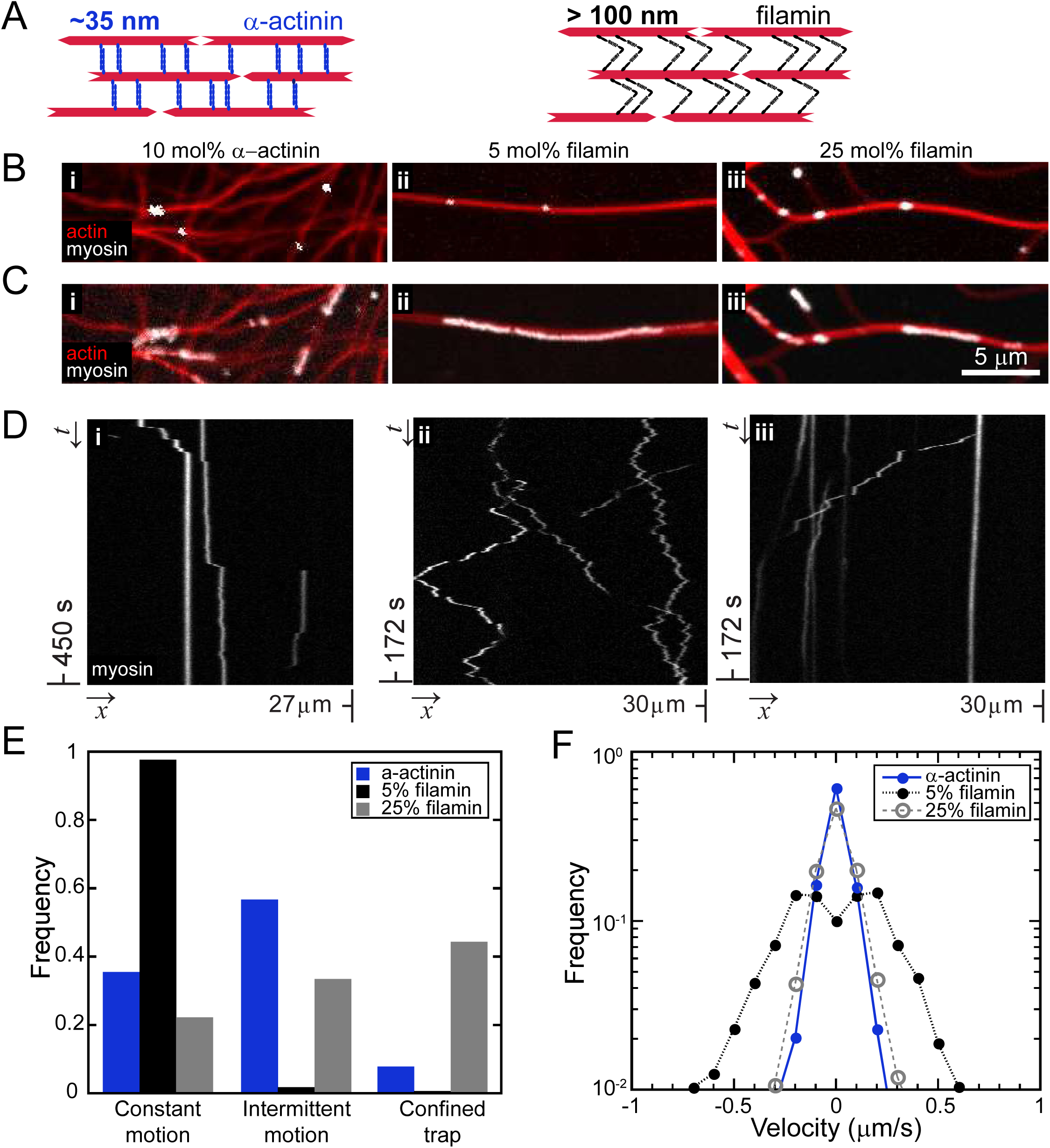
Bundle spacing and stiffness regulates myosin dynamics. (A) Cartoon of mixed polarity bundles constructed by the ∼35 nm, rigid cross-linker, α-actinin, and the ∼150 nm, hinged cross-linker filamin. (B) Bundles of F-actin (red) crosslinked by 10 mol% α-actinin (left), 5 mol% filamin (middle) and 25 mol% filamin (right) with isolated myosin II motors (white puncta). (C) Maximal intensity projection of myosin II on the bundles, showing that myosin II explores space on 5 mol% filamin bundles (middle), but on 10 mol% α-actinin (left) and 25 mol% filamin (right) bundles appears in localized, micron sized regions of the bundles (right). (D) Representative kymographs of myosin along different crosslinked bundles. 5 mol% filamin bundles (middle) show zig-zag diagonal lines with no vertical lines, indicating constant motion in the form of short runs in one direction, followed by runs in the opposite direction. Myosin II on 10 mol% α-acitnin (left) and 25 mol% filamin (right) bundles appears as long vertical lines interspersed with short, diagonal lines (middle), indicating motion dominated by trapping similar to fimbrin. (E) Summary of the frequency of myosin dynamics— intermittent motion and pauses, constant motion, or sustained pauses—from kymographs represented in (E). (α-actinin: 217, 5 mol% filamin: 167, 25 mol% filamin: 275 myosin) (F) Velocity distributions for the different crosslinked bundles are all symmetric about zero, indicating bidirectional motion. 5 mol% filamin (black dashed line and closed circles) has a minimum at zero, indicating little trapping and average non-zero speed ∼300 nm/s, while 10 mol% α-actinin (blue solid line and circle) and 25 mol% filamin (grey dashed line and open circles) have maxima at zero, indicating increased trapping.

To increase the interfilament spacing, we construct bundles using the cross-linker filamin, which has arms linked by disordered protein regions that result in a flexible hinge-like structure (Fig. 5A & 5Bii) (4, 42). Bundles constructed from F-actin cross-linked by 5 mol% filamin are loosely spaced collections of F-actin. In contrast to the smaller spaced bundles, projections of myosin over a 5 min period form tracks along several microns of filamin cross-linked bundles (Fig. 5Cii, white, Movie S6). Kymographs along filamin cross-linked bundles reveals markedly different myosin behavior, where tracks appear as zig-zagging lines, indicating direction switching over multiple length scales (Fig. 5Dii). Notably, in contrast to on the other mixed-polarity bundles, kymographs of myosin motion on filamin bundles contain no vertical lines of measurable duration, indicating that filamin bundles support continuous motion without confinement, with instantaneous velocity up to ∼500 nm/s (Fig. 5F, black dashed, closed circles). Intriguingly, increasing the number of filamin cross-links in a bundle to 25 mol% restores myosin motion to the motion characteristic of smaller spaced mixed polarity bundles (Fig. 4Biii & 4Ciii), where periods of motion are interspersed with traps (Fig 5Diii, Movie S7). The distribution of myosin that move continuously, intermittently, and are confined are similar to myosin on fimbrin bundles (Fig. 5E, grey, Fig. 2F, black). In the regions of motion, myosin on high filamin bundles has a speed ∼300 nm/s, consistent with myosin movements on fimbrin cross-linked actin (Fig. 5F, grey, open).

### Bundle spacing and compliance influence myosin trapping and force generation

To systematically investigate the influence of bundle spacing on myosin activity, we simulated a myosin filament on a mixed polarity bundle (*p* = 0.5, *N*_+_=*N*_-_=3, *k*=10 pN/nm) with variable interfilament spacing. Similar to our experimental observations, we find that myosin filaments, with *N*_heads_ =200, are trapped in sustained pauses at the smallest spacing, *s* = 10, but switch between intermittent runs and confined traps at the largest spacing, *s* = 200 nm (Fig. 6A).

**Figure 6:**
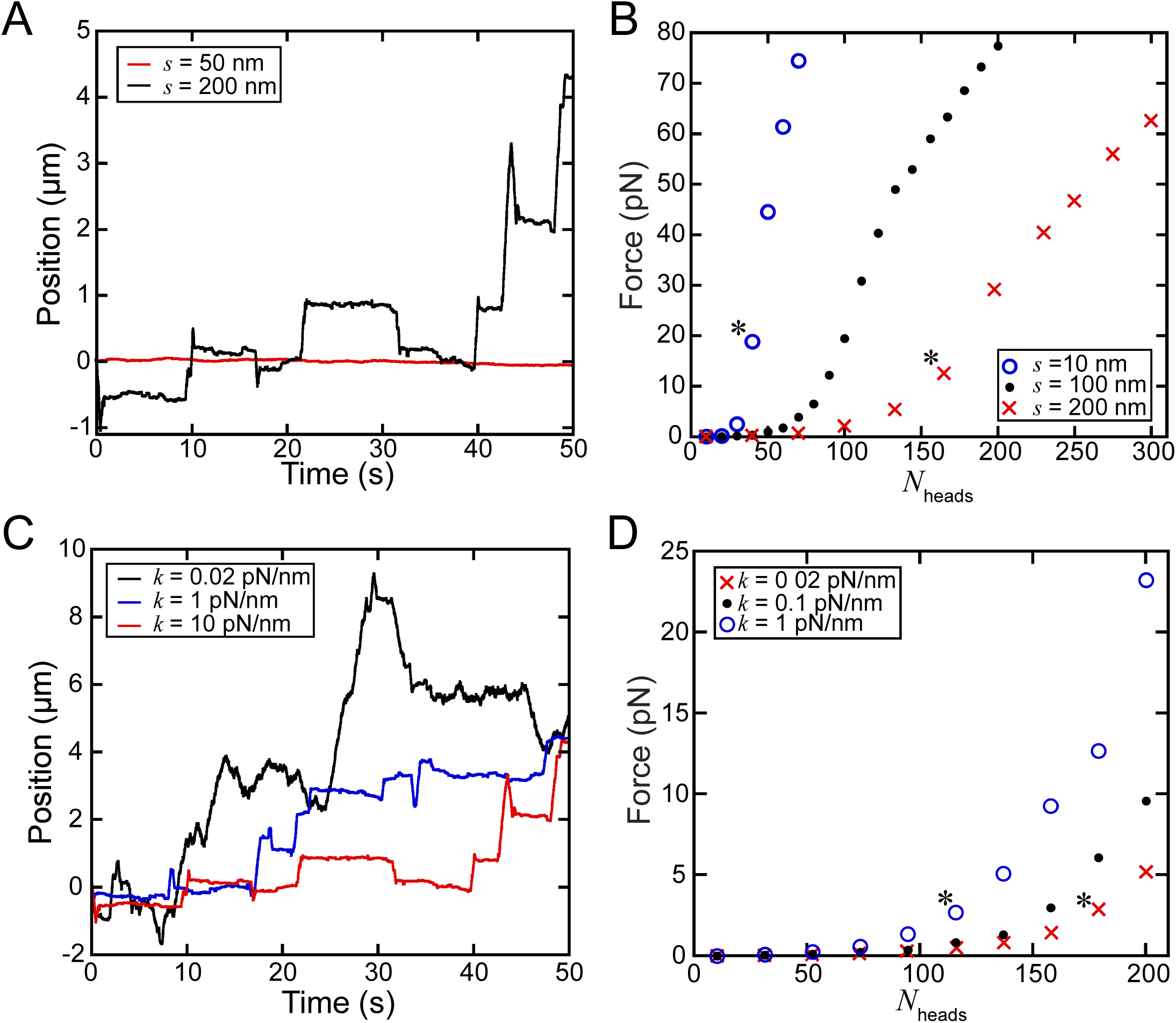
Forces exerted by myosin filaments at varying F-actin spacing cannot be readily inferred from velocity or distance travelled. (A) Trajectories on a bundle with *p* =0.5, *k* = 10 pN/nm, *N*_heads_ = 200, spacing *s* = 50 nm and 200 nm. (B) The number of motor domains, *N*_heads_, and spacing, *s*, affects the time-averaged force, *F*. At increasing *s*, indicated by darker grey, the transition to nonzero force production with increased *N*_heads_ is less steep. Value of *k* is 10 pN/nm. Asterisks indicate data in 6D. (C) Myosin filaments transition from weak attachment/noisy trajectories at low stiffness to tight trapping with increased stiffness. Representative trajectories at *s* = 200 nm, *N*_heads_ = 200. (D) Force output as a function of *N*_heads_, for different stiffness bundles with *s* = 200 nm. Asterisks indicate data in 6B.

To understand the impact of bundle spacing on forces generated by myosin, we quantified average force as a function of *N*_heads_. On a tightly spaced bundle, the force is near zero below a critical value (∼10 heads), and then increases linearly with increasing *N*_heads_ (Fig. 6B, blue), consistent with previous reports of myosin motors on individual F-actin (26). At or above the critical *N*_heads_ value, the force-dependent increase in the bound lifetime of myosin heads results in a positive feedback between force generation and myosin attachment (26). Below the critical number of myosin heads bound, the entire myosin filament frequently detaches, resulting in force relaxation, while above the myosin filament remains bound for long times (26). The critical *N*_heads_ to transition to increasing force increases with *s*, while the slope of the force dependence on *N*_heads_ decreases with *s* (Fig. 6B). The dependence of the force-*N*_heads_ relationship on interfilament spacing suggests that different spacings can support different force outputs for myosin with the same value of *N*_heads_.

In addition to differences in cross-linker size, which controls interfilament spacing, cross-linkers such as filamin and α-actinin differ in stiffness, which controls the bundle’s mechanical compliance. Decreasing the stiffness of the elastic load on which myosin motors build force decreases the total force output (26, 40, 43-45). This occurs because decreasing the stiffness reduces the rate of force buildup and reduces the positive feedback between myosin attachment time and force buildup (26, 40). Indeed, we find that for bundles with large spacing (*s* = 200, *p* = 0.5, *N*_+_=*N*_-_=3,) the motion exhibits complex behavior as a function of cross-linker stiffness (Fig. 6C). The myosin filaments are the most motile at the lowest value of stiffness, *k* = 0.02 pN/nm. This arises both from longer runs of the myosin filament in a given direction as well as differences in the nature of the trapped regions. At *k* = 0.02 pN/nm, there are larger position fluctuations in regions of the trajectory where the myosin filament is relatively trapped (e.g., Fig. 6C, black curve, 35-45 s) as compared to trapped regions at higher *k* in Fig. 6C. For a given *N*_heads_, the average force increases with increasing stiffness, while the critical *N*_heads_ for myosin to exert a non-zero force increases with decreasing stiffness (Fig. 6D). Together these data suggest that differences in both length and flexibility account for the lower forces produced by myosin on bundles formed by filamin.

We note that for a given force output myosin trajectories vary as a function of bundle spacing or stiffness, making direct inference of forces from these dynamics challenging. For stiff bundles (*k*=10 pN/nm, *F*=15-20 pN), corresponding to asterisks in Fig. 6B, myosin is completely trapped at the smallest spacing (*s* = 10 nm, Fig. 7A, red), but exhibits intermittent motion at the largest spacing (Fig. 7A, black). Similarly, we find that myosin with similar force outputs are characterized by very different motilities for bundles of different stiffness (Fig. 6B, corresponding to Fig. 6D, asterisks). For large spacing (*s*=200 nm), myosin is relatively trapped with intermittent short runs at extreme stiffnesses (Fig. 7B, black, *k* = 1 pN/nm), but moves in long runs frequented by direction switching at intermediate stiffness (Fig. 7B, red, *k* = 0.02 pN/nm).

**Figure 7.**
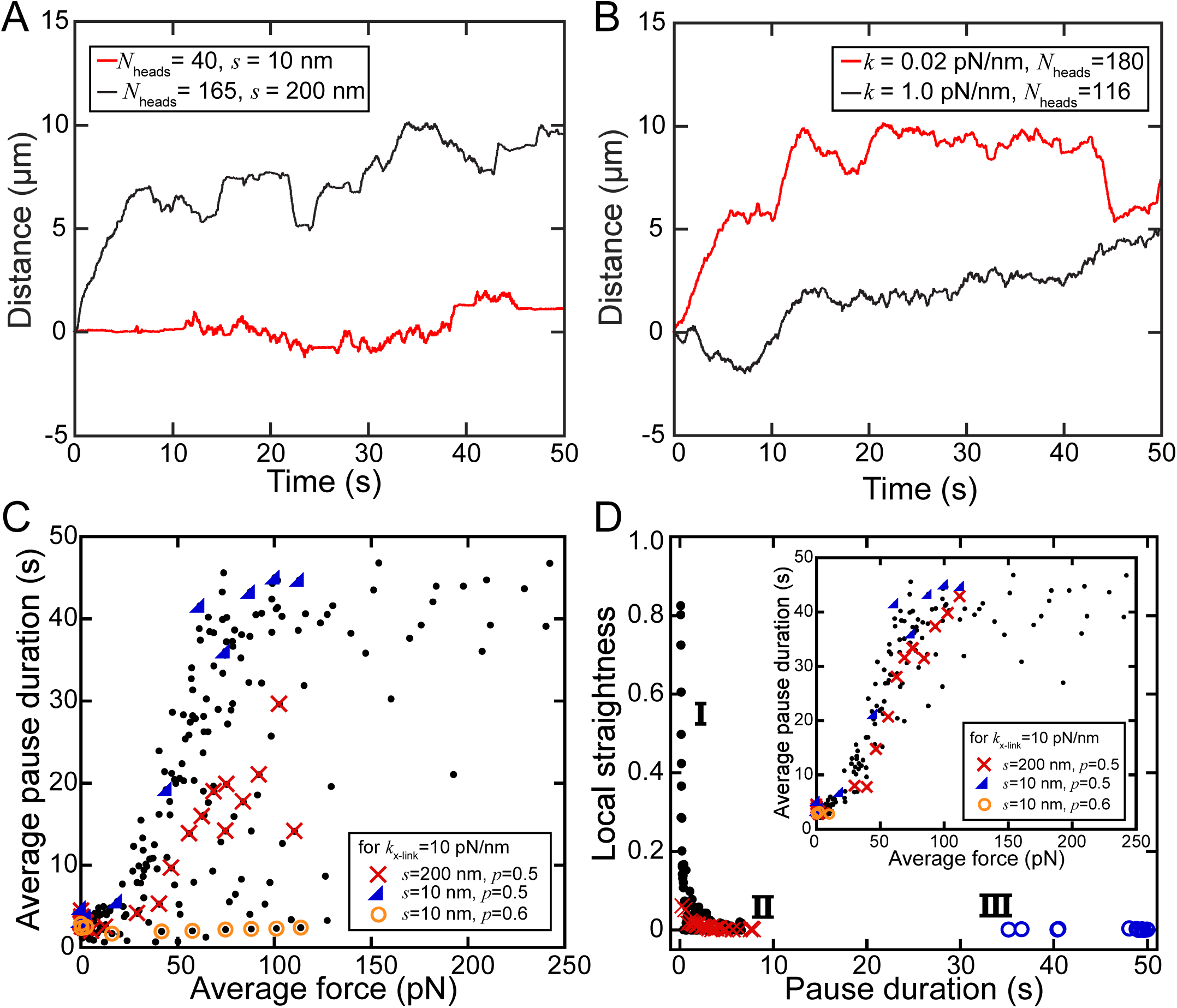
Pause time and trajectory straightness reveal qualitative differences in force generation. (A) Trajectories for myosin on bundle with different spacing, *s*=10 nm (red, *N*_heads_=40) and *s*=200 nm (black, *N*_heads_=165) at the same force output exhibit different motility characteristics. Force corresponds to data points from blue open circle and red x curves in Fig. 6B. (B) Trajectories for myosin at the same force output on different stiffness, *k* = 0.02 pN/nm (red, *N*_heads_=180) and *k* = 1 pN/nm (black, *N*_heads_=116), bundles exhibit different motility characteristics. Force corresponds to asterisks in Fig. 6B. (C) The average pause duration, defined as the height of peaks in the distribution of myosin position, plotted against average force for each set of parameters in Figs. 3, 4, and 6 including all values of *N*_heads_, *s, k*, and *p*. Highlighted examples are rigid bundles with small spacing (*k* = 10 pN/nm, *s* = 10 nm, *p* = 0.5, blue triangles), rigid bundles with larger spacing (*k* = 10 pN/nm, *s* = 200 nm, *p* = 0.5, red x’s) and rigid polar bundles (*k*_\_= 10 pN/nm, *s* = 10 nm, *p* = 0.6, orange circles). (D) Pause duration for each peak in the myosin position histogram against the local straightness. In region I, the pause duration is small because the myosin is continually moving on a straight trajectory. Myosin binds with high affinity but exerts variable forces in this regime with unloaded speeds indicating a lack of force production. In region II, the pause duration is small because myosin is weakly attached and diffuses. In region III, myosin is tightly trapped because it binds with high affinity to an apolar region and produces high forces. Data is from simulations with small actin spacing and stiff cross-linkers (*s* = 10 nm, *k* = 10 pN/nm) and *N*_heads_ = 20 (red x’s) or *N*_heads_ = 100 (blue circles). The black circles indicate data points corresponding to the trajectory in Fig. 6A (red) (*N*_heads_ = 200, *s* = 200, *k* = 10 pN/nm). (D, *inset*) Average pause duration quantified only for peaks in the myosin position distribution where the straightness parameter of the local myosin filament trajectory that is less than 0.1. When regions of locally straight myosin motion are excluded from the pause duration analysis in (C), the data collapses to a single curve.

### Trap strength is directly related to the average generated force

To extract information about myosin-generated forces from myosin trajectories, we detect traps by plotting the probability that a myosin filament is located in a specific position along the bundle during each 50 s simulation and find local peaks in the probability distribution. The height of each peak corresponds to the trap strength. We average the trap duration as well as the force during 10 independent 50 s simulations using systematically varied parameter values of *N*_heads_, bundle spacing, and cross-link stiffness (Fig. 7C). The average trap duration is highly variable with respect to the average force, with different bundle architectures dominating different regions of the pause space (Fig. 7C; e.g. blue triangles: *k* = 10 pN/nm, *p* = 0.5, *s* = 10 nm; red x’s: *k* = 10 pN/nm, *s* = 200 nm, *p* = 0.5; orange circles: *k* = 10 pN/nm, *s* = 10 nm, *p* = 0.6; each data point is for trajectories with one value of *N*_heads_). This variability is consistent with sensitive dependence of myosin force generation on the microscopic bundle architecture and motor binding affinity.

To further distinguish between high-force and low-force signatures in trajectories, we consider the trajectory straightness, defined as the ratio between the final displacement and the path length. Transient traps with small durations can occur either because the myosin is mostly unbound to actin, or because the myosin is able to bind but is continually walking in a direction determined by the polarity of its interactions with F-actin. In the former scenario, the force is very small due to the transient nature of the binding interactions; in the latter scenario, the force is variable and determined by the force-velocity relationship of the motors. These two situations can be distinguished from one another by plotting the strength of an individual trap against the trajectory straightness within a trap (Fig. 7D). Transient, low-force interactions such as those occurring at small *N*_heads_ (*N*_heads_ = 20, *s* = 10 nm *N*_+_=*N*_-_=3, Fig. 7D, region II) have relatively small values of both trap strength and trajectory straightness. On the other hand, myosin filaments that interact more strongly with the local F-actin but do so in a polar fashion display small values of trap strength but high values of straightness (*N*_heads_ = 200, s = 200 nm *N*_+_=*N*_-_=3, Fig. 7D, region I). Traps with long durations that are highly force-generating have small values of local trajectory straightness (*N*_heads_ = 100, s = 10 nm *N*_+_=*N*_-_=3, Fig. 7D, region III). Using this metric to differentiate between different traps, we discard traps with highly variable forces where the myosin filament moves with a straightness parameter >0.1, and find that the average force is directly proportional to pause duration for forces <100 pN (Fig. 7D, *inset*). This indicates that myosin in weak traps produce little or no forces, while myosin in strong traps generate large forces.

## Discussion

Cells have a myriad of cytoskeletal architectures which mediate physiological processes as diverse as directing transport to supporting cellular shape change. Here, we find that diverse F-actin bundle architectures constructed from physiological cross-linkers support different dynamics of myosin II complexes. Bundle architecture has been shown to influence protein segregation (46). Steric considerations such as interfilament spacing can influence protein binding, as has been shown for cross-linker binding (10, 11). Similarly, filament polarity and interfilament spacing within bundles has been shown to influence motor transport *in vitro* (12, 16, 47). Regional differences in bundle architecture have been shown to influence the localization of proteins, such as different myosin isoforms, to different areas of the cell (17, 48). Future research might explore the further contributions of differences in interfilament angle, such as those observed in branched actin networks assembled by ARP 2/3.

While architecture has been hypothesized to regulate protein segregation and cellular transport due to motor preference, the effects of architecture on cellular force generation is a nascent field. Mixed polarity architecture in bundles has been shown to promote contractility (49), and the deformations depend on actin bundling (20). Here, we relate the experimentally observed motor dynamics to force generation through agent-based simulations. Our research has focused on filaments of skeletal muscle myosin II, which forms larger motor complexes than the isoforms of myosin II found in non-muscle cells. Since our simulations suggest that the force-generation potential depends sensitively on the number of motor heads in a complex, it would be interesting to explore how different isoforms of myosin II respond to different actin architectures.

Our results have implications for how force generation may be spatially regulated in the cell through bundle mechanics. Intriguingly, we show that compliant, filamin cross-linked bundles can be tuned, switching from negligible force production (loose traps) to tight traps or runs with increased cross-linker stiffness. Crosslinkers and other F-actin binding proteins could potentially dynamically tune local bundle mechanics, leading to spatial and temporal control of force generation. The different dependencies of myosin isoforms on mechanical feedback in building force (26, 50) might cause bundle mechanics to shape the force response to different isoforms. We hypothesize that the isoform myosin IIa will be the most sensitive to bundle mechanics due to its force building mechanism. These and other isoform-specific properties may play a role in differential localization of non-muscle myosin II isoforms observed within cells (51). Together, this work presents a framework for the analysis of force generation in cytoskeletal networks inside and outside of cells. Additionally, understanding the effects of microstructure on force generation has the potential to inform the design of artificial active polymeric materials with tunable force response.

## Acknowledgements

We thank members of David Kovar Laboratory for purified protein cross-linkers, particularly J. Winkelman, C. Suarez, J. Christensen, and Y. Li. This research was partially supported by the University of Chicago MRSEC, funded through the NSF award # DMR-1420709. This research was funded by the NSF DMR-1905675 and MCB-1344203 to M.L.G., the National Institute of Genemral Medical Sciences grant # 1R01GM098441 to E.M.M., and the NIH National Institute of Biomedical Imaging and Bioengineering Training grant # T32EB009412 (S.S.)

